# Tau controls NMDA receptor trafficking during homeostatic synaptic plasticity

**DOI:** 10.1101/2025.07.28.667139

**Authors:** Xuan Ling Hilary Yong, Kristie Stefanoska, William C. Kwan, Isabelle Irons, Pranesh Padmanabhan, Evgeniia Samokhina, Andrew Kneynsberg, Nathalie Dehorter, Arne Ittner, Jürgen Götz, Victor Anggono

## Abstract

Homeostatic synaptic plasticity is essential for maintaining stable neural circuit function by preventing excessive neuronal excitation or inhibition. Chronic perturbation of neuronal activity triggers a compensatory modulation of the number of α-amino-3-hydroxy-5-methyl-4-isoxazole propionic acid (AMPA) and *N*-methyl-*D*-aspartate (NMDA) glutamate receptors at the excitatory synapses. Previous research has primarily focused on AMPA receptors, yet the molecular mechanisms regulating the trafficking of NMDA receptors during homeostatic synaptic scaling remain unclear. Here we identify the microtubule-associated protein Tau as an essential molecule that mediates the synaptic upscaling of GluN2B-containing NMDA receptors during prolonged synaptic inactivity. Chronic activity blockade increases Tau phosphorylation at Ser-235 by cyclin-dependent kinase 5 (Cdk5), enhancing its interaction with and retention of active Fyn tyrosine kinase in the postsynaptic compartment. This promotes the phosphorylation of GluN2B at Tyr-1472, subsequently stabilising the expression of NMDA receptors on the neuronal plasma membrane. Finally, we showed that Tau pathology and disease-associated mutations in Tau and the GluN2B carboxyl-terminal tail disrupt the homeostatic synaptic upscaling of NMDA receptors following chronic neuronal silencing. Together, our findings identify a physiological role for Tau in homeostatic synaptic plasticity, the perturbation of which can lead to neuronal hyperexcitation, seizures and excitotoxic cell death.

## INTRODUCTION

The interplay between Hebbian and homeostasis synaptic plasticity maintains neuronal excitation-inhibition balance within the neural circuits to support learning and memory formation (Keck et al., 2017; Turrigiano, 2017). Hebbian forms of plasticity, such as long-term potentiation (LTP) and long-term depression (LTD), drive persistent changes in synaptic strength and, when uncorrected, can lead to runaway excitation or inhibition (Turrigiano, 2008; Caya-Bissonnette and Béïque, 2024). Hence, homeostatic processes are necessary to counteract these effects by stabilising neuronal firing rate at an optimal set point (Turrigiano, 1999; Davis and Bezprozvanny, 2001; Marder and Prinz, 2003; Turrigiano and Nelson, 2004). One form of homeostatic plasticity, termed synaptic scaling, provides a cell-wide, bidirectional and proportional adjustment in the number and composition of receptors and ion channels at synapses to fine-tune neuronal excitability (O’Brien et al., 1998; Turrigiano et al., 1998). Homeostatic synaptic scaling has been observed in sensory cortices *in vivo* (Maffei et al., 2004; Goel et al., 2006; Keck et al., 2013), and plays a role in memory formation (Wu et al., 2021). Perturbations of neuronal network homeostasis have been implicated in the early pathogenesis of Alzheimer’s disease (AD) (Frere and Slutsky, 2018; Styr and Slutsky, 2018) and can also cause neuronal hyperactivity, commonly observed in patients with AD (Palop and Mucke, 2010).

NMDA receptors are the primary source of calcium (Ca^2+^) at the postsynaptic compartment and are composed of two obligatory GluN1 subunits and two identical (diheteromeric) or different (triheteromeric) GluN2 subunits (Paoletti et al., 2013; Vieira et al., 2020). The glutamate-binding GluN2 subunits are encoded by four genes (*GRIN2A* – *GRIN2D*) in the mammalian brain, each of which confers NMDA receptors with distinct ion channel properties and intracellular trafficking pathways (Vieira et al., 2020; Hansen et al., 2021). GluN2B-containing NMDA receptors have a lower channel opening probability and a slower deactivation time than those containing GluN2A, such that an activity-dependent switch in NMDA receptor subunit composition at synapses has major implications for synaptic integration, network synchrony and synaptic plasticity (Yashiro and Philpot, 2008). GluN2B-containing NMDA receptors are essential for normal brain development, synaptic plasticity, learning and memory (von Engelhardt et al., 2008; Brigman et al., 2010). However, aberrant over-activation of GluN2B can lead to intracellular Ca^2+^ overload, neuronal hyperexcitation (seizures) and excitotoxicity, a phenomenon tightly linked to neurodegenerative diseases, including AD (Hardingham and Bading, 2010). Therefore, the number of surface GluN2B-NMDA receptors must be tightly regulated to allow normal brain function without causing neuronal death.

In response to altered network activity, neurons bidirectionally scale the number of α-amino-3-hydroxy-5-methyl-4-isoxazole propionic acid (AMPA) and *N*-methyl-*D*-aspartate (NMDA) glutamate receptors at the excitatory synapses (Watt et al., 2000; Soares et al., 2013). Unlike AMPA receptors, the molecular mechanisms that underpin the homeostatic synaptic scaling of NMDA receptors remain unknown. In this study, we focused on the microtubule-associated protein Tau, best known for its disease-enabling roles when it is mislocalised in the dendritic compartment, hyperphosphorylated and/or aggregated in several neurodegenerative diseases (Ittner and Ittner, 2018; Polanco et al., 2018; Chang et al., 2021a). In AD and stroke models, GluN2B-dependent excitotoxic signalling is exacerbated by the mislocalisation of Tau to dendritic spines (Ittner et al., 2010; Bi et al., 2017). Although Tau is enriched in the axons, increasing evidence indicates that a small fraction of Tau is also present in the somatodendritic compartment and dendritic spines under physiological conditions (Chang et al., 2021a; Parra Bravo et al., 2024). Here, we reveal that Tau plays a critical role in regulating the surface expression of NMDA receptors during homeostatic synaptic scaling, uncovering a physiological (non-pathological) function of Tau in maintaining glutamate receptor homeostasis in mammalian central neurons.

## RESULTS

### Tau mediates homeostatic synaptic upscaling of GluN2B-NMDA receptors

Homeostatic synaptic scaling can be modelled in primary hippocampal neurons by treating them with the voltage-gated sodium channel blocker tetrodotoxin (TTX) for 24 h (Anggono et al., 2011). We first performed surface biotinylation assays and found that chronic silencing induced by TTX treatment resulted in a robust compensatory increase in the surface expression of GluN2B-, but not GluN2A-containing NMDA receptors, in wild-type primary hippocampal neurons (Figure 1A and 1B, Supplementary Figure 1), confirming the findings from previous studies (Ehlers, 2003; Bateup et al., 2013). This effect was abolished in Tau knockout (Figure 1A and 1B) and Tau knockdown neurons (Supplementary Figure 2A and 2B). Interestingly, chronic TTX treatment caused a significant reduction in the level of total Tau protein in wild-type neurons (Figure 1A and 1C). These results demonstrate not only an essential physiological role of Tau in controlling the homeostasis of surface NMDA receptor expression in a subunit-dependent manner, but also reveal that the Tau protein abundance is regulated by neuronal activity.

**Figure 1.**
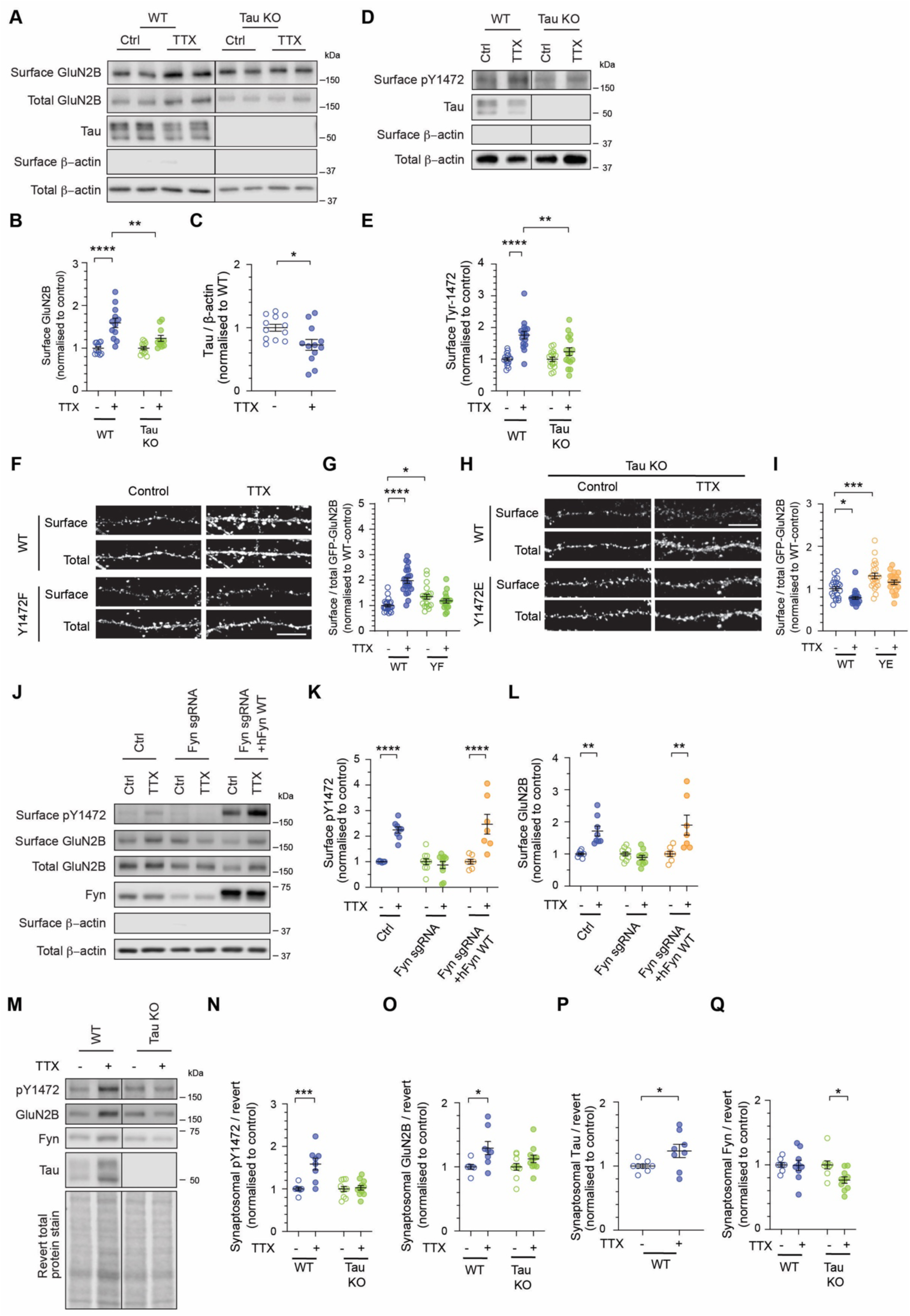
Tau mediates Fyn-dependent phosphorylation of GluN2B on Tyr-1472 during homeostatic synaptic scaling. (A) Primary hippocampal neurons derived from wild-type and Tau knockout mice were treated with water (control) or 2 μM tetrodotoxin (TTX) for 24 h. The relative amounts of surface and total proteins were assessed by western blotting using specific antibodies against GluN2B, Tau and β-actin. Tau blot confirmed the genotype of cultured neurons. (B) Quantification of the surface GluN2B, normalised to the control group (n = 12 cultures per group from 3 independent experiments). ** P < 0.01, **** P < 0.0001 using two-way ANOVA with a Tukey’s multiple comparisons test. All data represent mean ± SEM. (C) Wild-type neurons treated with TTX result in a reduction in total Tau protein expression (n = 12 cultures per group from 3 independent experiments). * P < 0.05 using an unpaired Mann-Whitney t test. All data are represented as mean ± SEM. (D) Wild-type and Tau knockout primary hippocampal neurons were treated with water (control) or 2 μM TTX for 24 h and subjected to a surface biotinylation assay. The relative amounts of surface and total proteins were assessed by immunoblotting with specific antibodies against phosphorylated Tyr-1472, total Tau, and β-actin. Tau blot confirmed the genotype of cultured neurons. (E) Quantification of the surface Tyr-1472 phosphorylation levels normalised to the control group (n = 9-16 cultures per group from 4 independent experiments). ** P < 0.01, **** P < 0.0001 using two-way ANOVA with a Tukey’s multiple comparisons test. All data represent mean ± SEM. (F) Wild-type hippocampal neurons expressing either GFP-tagged GluN2B-wild-type or Tyr-1472F phospho-deficient mutant were stimulated with TTX for 24 h. Live neurons were incubated with a rabbit anti-GFP antibody at room temperature to label surface GFP-tagged GluN2B, followed by fixation, permeabilisation, and staining with a mouse anti-GFP antibody to detect total GFP expression. Representative enlarged images of dendritic segments showing the surface and total GFP staining for each group. Scale bar, 5 μm. (G) Quantification of the surface/total GFP ratio normalised to that of non-stimulated neurons (n = 17– 23 neurons per group from 2 independent experiments). (H) Tau KO hippocampal neurons expressing either GFP-tagged GluN2B-wild-type or Tyr-1472E phospho-mimetic mutant were stimulated with TTX for 24 h. Representative enlarged images of dendritic segments showing the surface and total GFP staining for each group. Scale bar, 5 μm. (I) Quantification of the surface/total GFP ratio normalised to that of non-stimulated neurons (n = 20– 23 neurons per group from 2 independent experiments). ** P < 0.01, *** P < 0.001, **** P < 0.0001 using two-way ANOVA with a Tukey’s multiple comparisons test. All data represent mean ± SEM. (J) Primary hippocampal neurons were transduced with lentiviral particles expressing Cas9-GFP and Fyn sgRNA, at DIV9. At DIV13, Fyn knockdown neurons were further transduced with lentiviral particles expressing Fyn-wild-type-Myc. Water control or TTX-treated neurons were subjected to a surface biotinylation assay after 24 h treatment. The relative amounts of surface and total proteins were assessed by immunoblotting with specific antibodies against Tyr-1472, GluN2B, Fyn, Tau, and β-actin. (K-L) Quantification of the surface Tyr-1472 phosphorylation levels (K), and GluN2B surface levels normalised to unstimulated groups (n = 7–11 cultures per group from 3 independent experiments). ** P < 0.01, **** P < 0.0001 using two-way ANOVA with a Tukey’s multiple comparisons test. All data represent mean ± SEM. (M) After 24 h TTX treatment, cultured hippocampal neurons derived from wild-type and Tau knockout mice were subjected to subcellular fractionation to isolate the synaptosome fractions. The relative amounts of proteins were assessed by immunoblotting with specific antibodies against Tyr-1472, GluN2B, Fyn, Tau. Revert total protein stain was used to visualise overall protein content. (Ν-Q) Quantification of Tyr-1472 phosphorylation (N), and GluN2B (O), Tau (P) and Fyn (Q) levels normalised to Revert. Data is expressed relative to the unstimulated groups (n = 7–11 cultures per group from 3 independent experiments). *** P < 0.001, using two-way ANOVA with a Tukey’s multiple comparisons test, except for Tau, which was analysed using an unpaired Mann-Whitney t test * P < 0.05. All data represent mean ± SEM.

### Tau-dependent phosphorylation of GluN2B on Tyr-1472 is required for synaptic upscaling of NMDA receptors

One of the key determinants that controls the trafficking and function of NMDA receptors is the phosphorylation at the carboxyl-terminal domain of GluN2 subunits (Lussier et al., 2015; Vieira et al., 2020). Two major phosphorylation sites on GluN2B carboxyl-terminal tail, Tyr-1472 and Ser-1480, are known to regulate the surface expression and internalisation of NMDA receptors (Lavezzari et al., 2003; Chung et al., 2004; Prybylowski et al., 2005; Sanz-Clemente et al., 2010). Consistent with previous findings, prolonged neuronal silencing induced a robust increase in the level of Tyr-1472 phosphorylation and markedly reduced Ser-1480 phosphorylation on the GluN2B subunit (Figure 1D and 1E, Supplementary Figure 3A and 3B) (Sanz-Clemente et al., 2010; Jang et al., 2015). Interestingly, TTX-induced upregulation of Tyr-1472 phosphorylation was abolished in Tau knockout neurons (Figure 2D and 2E). Conversely, TTX-induced dephosphorylation of Ser-1480 remained intact in Tau knockout neurons (Supplementary Figure 3A and 3B). We also confirmed these findings in single guide RNA (sgRNA)-mediated Tau knockdown neurons (Supplementary Figure 2A, 2C and 2D). These data indicate that Tau-dependent phosphorylation of Tyr-1472, which prevents the binding of endocytic adaptor AP-2 to GluN2B (Lavezzari et al., 2003; Prybylowski et al., 2005), underpins the increase in surface NMDA receptor expression during TTX-induced synaptic scaling.

**Figure 2.**
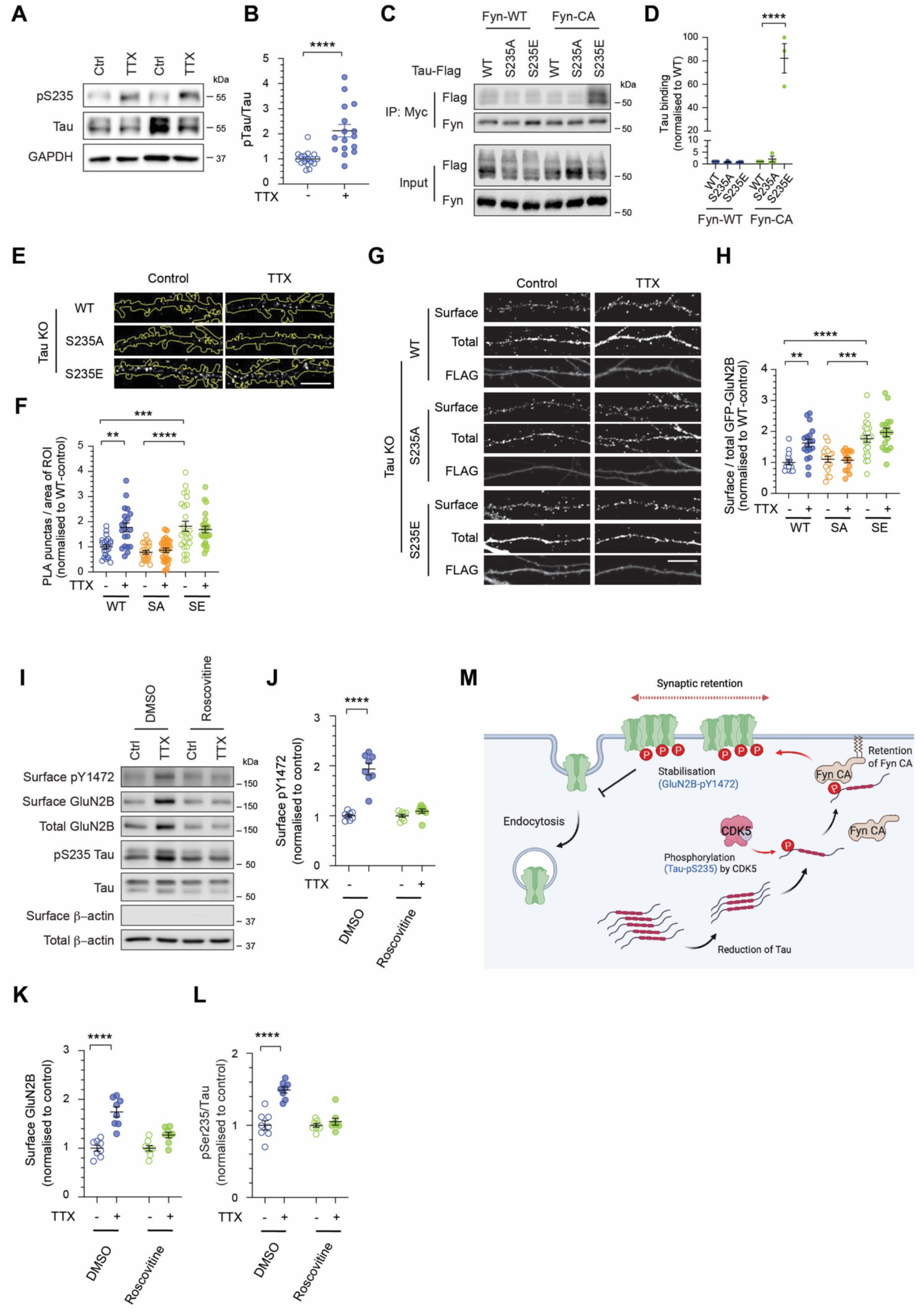
Tau phosphorylation on Ser-235 by Cdk5 is required for TTX-induced synaptic upscaling of GluN2B-NMDA receptors. (A) Total sample lysates were prepared from wild-type hippocampal neurons following 24 h water control or TTX treatment. The relative amounts of proteins were assessed by western blotting using antibodies specific for phosphorylated Tau at Ser-235, total Tau, and GAPDH. (B) Quantification of the ratios of Ser-235/Tau (n = 16 cultures per group from 4 independent experiments). All data are represented as mean ± SEM. **** P < 0.0001 using an unpaired Mann-Whitney t test. (C) HEK293T cells were co-transfected with plasmids encoding Fyn-Myc (wild-type or constitutively active; CA) and either hTau-Flag wild-type, S235A or S235E. HEK cell lysates were immunoprecipitated (IP) with antibodies against Myc. Total neuronal lysates (Input) and co-immunoprecipitated proteins were analysed by immunoblotting with specific antibodies against Flag and Fyn. (D) Quantification of the levels of Fyn binding to Tau. Data represent mean ± SEM of band intensities normalised to hTau-Flag wild-type (n = 3). **** P < 0.0001 using one-way ANOVA with Tukey’s multiple comparison test. (E) Tau KO hippocampal neurons expressing either hTau-Flag wild-type, S235A or S235E were stimulated with TTX for 24 h and subjected to a proximity ligation assay (PLA) with Fyn and Flag antibodies. Representative images showing the PLA signals in the dendrites. Scale bars, 10 μm. (F) Quantification of the number of PLA puncta in the dendrites. Data are normalised to the unstimulated group (n = 23–26 neurons per group from 4 independent experiments). All data are represented as mean ± SEM. ** P < 0.01, *** P < 0.001, **** P < 0.0001 using one-way ANOVA with Tukey’s multiple comparison test. (G) Tau KO hippocampal neurons expressing GFP-tagged GluN2B-wild-type with either hTau-Flag wild-type, S235A or S235E were stimulated with TTX for 24 h. Live neurons were incubated with a rabbit anti-GFP antibody at room temperature to label surface GFP-tagged GluN2B, followed by fixation, permeabilisation, and staining with a mouse anti-GFP antibody to detect total GFP expression. Representative enlarged images of dendritic segments showing the surface and total GFP staining for each group. Scale bar, 5 μm. (H) Quantification of the surface/total GFP ratio normalised to that of non-stimulated neurons (n = 14– 24 neurons per group from 3 independent experiments). ** P < 0.01, **** P < 0.0001 using two-way ANOVA with a Tukey’s multiple comparisons test. All data represent mean ± SEM. (I) Primary hippocampal neurons were treated with either DMSO or 25 μM roscovitine, and within each treatment group, cells were further treated with water (control) or TTX for 24 hours and subjected to a surface biotinylation assay. The relative amounts of surface and total proteins were assessed by western blotting using specific antibodies against phosphorylated Tyr-1472, Ser-235, total GluN2B, Tau and β-actin. (J-L) Quantification of the surface Tyr-1472 phosphorylation levels (J), GluN2B surface (K) and Ser-235/Tau level normalised to unstimulated groups (n = 8 cultures per group from 3 independent experiments). **** P < 0.0001 using two-way ANOVA with a Dunnett’s multiple comparisons test. All data represent mean ± SEM. (M) A proposed model for Tau in regulating the trafficking of GluN2B-NMDARs during homeostatic plasticity. During TTX-induced homeostatic plasticity, CDK5 phosphorylates Tau at Ser-235 to promote the retention of activated tyrosine kinase Fyn at the postsynaptic compartment. This position Fyn near the GluN2B-containing NMDA receptors, enabling the phosphorylation of the GluN2B carboxyl-terminal tail at Tyr-1472.

To confirm the role of Tyr-1472 phosphorylation during TTX-induced synaptic upscaling of GluN2B, we first expressed GFP-tagged GluN2B (wild-type or the Y1472F phospho-deficient mutant) in wild-type neurons and performed the surface staining assay with anti-GFP antibodies that recognise the extracellular GFP on the GluN2B subunit. TTX-induced synaptic scaling led to a robust increase in the surface expression of GFP-GluN2B wild-type in primary hippocampal neurons (Figure 1F and 1G). In contrast, neurons expressing the Y1472F phospho-deficient mutant did not exhibit an increase in GluN2B surface expression upon TTX treatment (Figure 1F and 1G). These results demonstrate that Tyr-1472 phosphorylation on GluN2B is necessary for homeostatic synaptic upscaling of NMDA receptors. Next, we expressed GFP-GluN2B (wild-type or the Y1472E phosphor-mimetic mutant) in Tau knockout neurons and performed the same surface staining assay. Consistent with our surface biotinylation data, synaptic upscaling of GFP-GluN2B was abolished in Tau knockout neurons (Figure 1H and 1I). Importantly, the expression of pseudo-phosphorylated GFP-GluN2B Y1472E mutant in Tau knockout neurons led to a significant increase in surface GFP-GluN2B expression under basal conditions and upon TTX treatment (Figure 1H and 1I). The expression of GFP-GluN2B Y1472E in wild-type neurons also led to a significant increase in surface GluN2B expression, which occluded any further upregulation of NMDA receptors on the neuronal plasma membrane following chronic neuronal silencing (Supplementary Figure 4). Together, these results establish the functional significance of Tau in mediating the phosphorylation of Tyr-1472, which is necessary and sufficient to induce the upregulation of NMDA receptors during homeostatic synaptic upscaling.

### Tau controls Fyn-mediated Tyr-1472 phosphorylation during synaptic scaling

Fyn is the primary tyrosine kinase that phosphorylates GluN2B on Tyr-1472 (Nakazawa et al., 2001). Next, we tested the role of Fyn in mediating GluN2B phosphorylation upon prolonged neuronal inactivity. We found that sgRNA-mediated knockdown of Fyn in primary hippocampal neurons completely abolished TTX-induced upregulation of Tyr-1472 phosphorylation and surface GluN2B-NMDA receptors (Figure 1J-1L). These deficits were fully rescued by co-expressing human Fyn, the cDNA of which is resistant to mouse-specific Fyn sgRNA (Figure 1J-1L). Our results demonstrate that Fyn is the kinase that mediates Tyr-1472 phosphorylation and is required for TTX-induced synaptic upscaling of GluN2B-NMDA receptors.

Tau interacts with Fyn and regulates its subcellular localisation in neurons (Lee et al., 1998; Ittner et al., 2010; Padmanabhan et al., 2019). To investigate the influence of Tau on Fyn subcellular distribution during homeostatic synaptic scaling, we performed subcellular fractionation from wild-type and Tau knockout neurons following TTX treatment. Prolonged neuronal inactivity increased Tyr-1472 phosphorylation and the number of GluN2B-NMDA receptors in the synaptosomal fraction of wild-type neurons, which were absent in Tau knockout neurons (Figure 1M-1O). Although we observed a significant reduction of total Tau protein level upon TTX treatment (Figure 1A and 1C), there was a significant accumulation of Tau protein in the synaptosomes (Figure 1M and 1P). Intriguingly, TTX treatment did not result in an apparent change in the level of Fyn protein in the synaptosomes; however, we found a marked reduction of synaptosomal Fyn in Tau knockout neurons (Figure 1M and 1Q). These data indicate that during homeostatic synaptic scaling, Tau functions to retain Fyn kinase in the postsynaptic compartment to phosphorylate GluN2B on Tyr-1472, thereby stabilising the pool of surface NMDA receptor expression.

### Cdk5-dependent phosphorylation of Tau on Ser-235 mediates homeostatic upscaling of GluN2B-NMDA receptors

To understand the mechanism that regulates Tau-Fyn interaction during homeostatic synaptic scaling, we sought to identify Tau phosphorylation sites that are modulated in response to TTX treatment. We screen a panel of ten site-specific anti-phospho-Tau antibodies that span from the proline-rich domain, microtubule-binding region and the carboxyl-terminal domain by western blotting analysis. Among these phosphorylation sites, Ser-235 phosphorylation showed the most robust increase (Figure 2A and 2B). We also observed a small but significant increase in the level of Ser-396 phosphorylation following TTX treatment (Supplementary Figure 5). Given that the Ser-235 residue lies within the vicinity of the Fyn binding site (Lee et al., 1998; Lau et al., 2016), we hypothesise that phosphorylation of Ser-235 modulates the binding affinity of Tau to Fyn kinase. To test this, we co-expressed Flag-hTAU (wild-type, S235A or S235E) with Fyn-myc, either wild-type or the constitutively active Y531F mutant, in HEK293T cells and performed an immunoprecipitation assay. Our results showed a comparable binding between wild-type Fyn to wild-type Tau or the Ser-235A mutants (Figure 2C and 2D). However, the phosphomimetic Tau S235E mutant displayed a higher binding affinity to the constitutively active Fyn (Figure 2C and 2D), suggesting that Ser-235 phosphorylation enhances the interaction between Tau and active Fyn. To confirm this interaction in neurons, we transfected plasmids that encode Flag-hTAU (either wild-type, S235A or S235E mutants) in Tau knockout neurons and performed proximity ligation assay (PLA) using antibodies that recognise Flag and endogenous Fyn. We found that chronic neuronal silencing increased the number of PLA puncta between Flag-TAU wild-type and endogenous Fyn in the dendrites (Figure 2E and 2F). TTX-induced increase of hTAU-Fyn PLA puncta was inhibited by the expression of the S235A phospho-deficient, whereas neurons expressing the S235E phospho-mimetic mutant exhibited elevated levels of PLA puncta under basal conditions, which were not upregulated further upon synaptic upscaling (Figure 2E and 2F). These data indicate that the phosphorylation of Tau on Ser-235 mediates the TTX-induced upregulation of Tau-Fyn interaction in the postsynaptic compartment during homeostatic synaptic plasticity.

To determine the functional significance of this enhanced Tau-Fyn interaction on the synaptic upscaling of GluN2B-NMDA receptors, we examined the levels of surface GluN2B expression levels in Tau knockout neurons expressing hTAU (either wild-type, S235A or S235E mutants). As expected, re-expression of wild-type hTAU rescued the homeostatic synaptic upscaling of GluN2B-NMDA receptors in Tau knockout neurons, whereas the S235A phospho-deficient mutant failed to do so (Figure 2G and 2H). In contrast, the phospho-mimetic S235E mutant augmented GluN2B-NMDAR surface expression with or without TTX treatment (Figure 2G and 2H). Together, our results demonstrate that TTX-induced phosphorylation of Ser-235 is necessary and sufficient to enhance the surface expression of GluN2B during homeostatic synaptic scaling by promoting the interaction of Tau with the Fyn kinase in the dendrites.

Cyclin-dependent kinase 5 (Cdk5) is one of the protein kinases known to phosphorylate Tau on Ser-235 (Liu et al., 2002; Sakaue et al., 2005). To determine if Cdk5 is responsible for the increase of Ser-235 phosphorylation during homeostatic synaptic scaling, we acutely inhibited Cdk5 with roscovitine. Roscovitine treatment blocked TTX-induced increase in the phosphorylation of Tau at Ser-235 and consequently abolished the upregulation of Tyr-1472 phosphorylation and GluN2B surface expression in primary hippocampal neurons (Figure 2I-2L). Collectively, these data demonstrate that homeostatic synaptic upscaling of NMDA receptors is driven by Cdk5-mediated phosphorylation of Tau on Ser-235, leading to an enhanced interaction with activated Fyn kinase within the dendritic compartments, which in turn promotes GluN2B phosphorylation at Tyr-1472 and stabilises the surface expression of NMDA receptors on the neuronal plasma membrane (Figure 2M).

### Tau pathology and disease-associated Tau and GluN2B mutations impair homeostatic synaptic scaling of NMDA receptors

Aberrant homeostatic synaptic scaling contributes to the pathogenesis of many neurological disorders, including AD, epilepsy and stroke (Fernandes and Carvalho, 2016; Styr and Slutsky, 2018). Since the abnormal accumulation of intracellular Tau aggregates is a prominent pathological feature of AD, we next investigated if homeostatic synaptic scaling of NMDA receptors was affected by the presence of aggregation-prone TAU variants. We co-expressed GFP-GluN2B and various Flag-hTAU mutants in Tau knockout neurons and found that Tau P301L, AT8 (S202E/T205E/S208E) and 12E8 (S262E/S356E) caused a dramatic increase in the levels of surface GluN2B under basal conditions and impaired TTX-induced synaptic upscaling of surface NMDARs (Figure 3A and 3B). These data indicate that these aggregation-prone TAU variants are toxic gain-of-function mutants that cause aberrant dysregulation of NMDA receptor surface expression by impairing the homeostatic synaptic plasticity mechanism in neurons.

**Figure 3.**
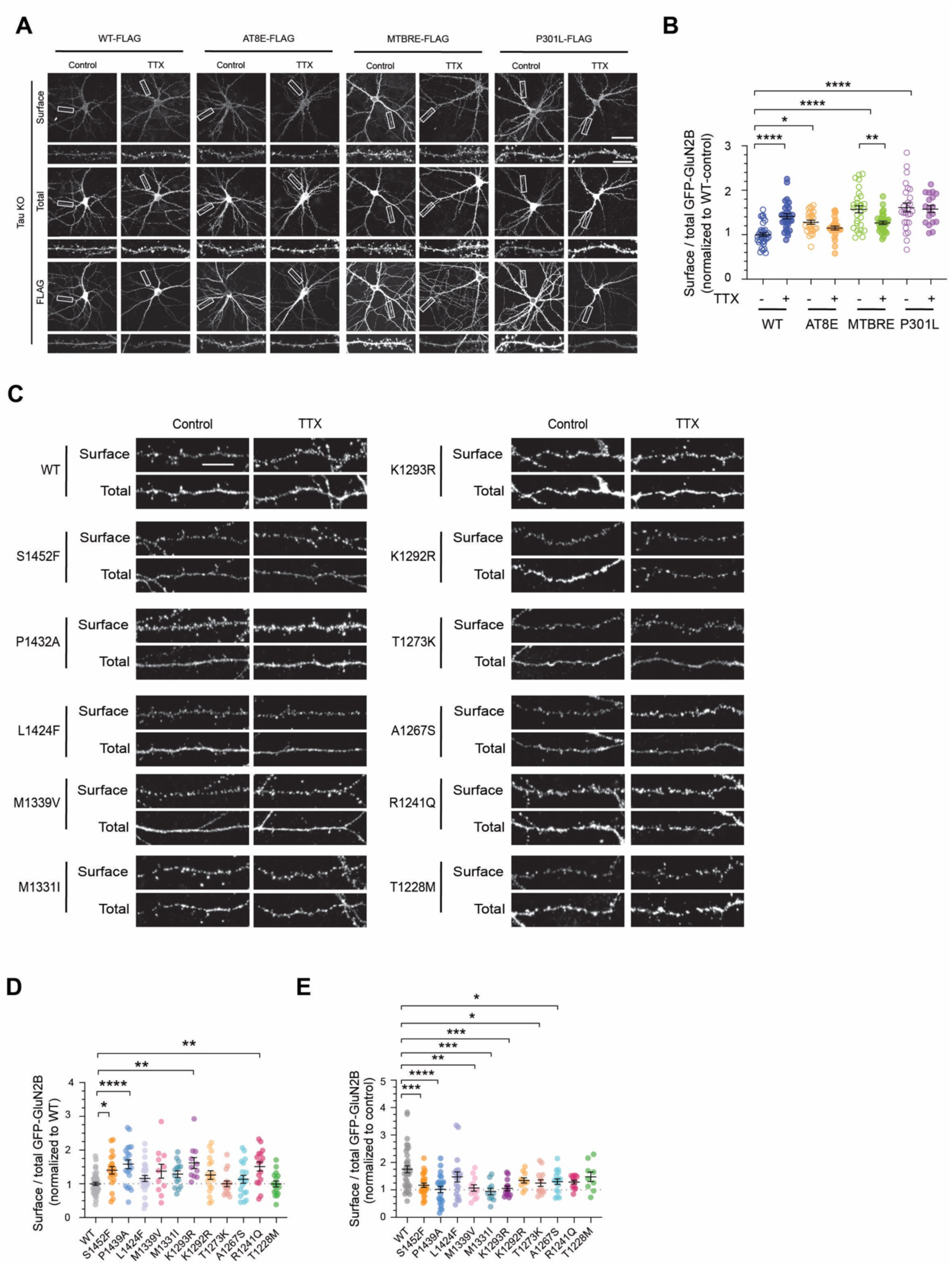
Pathogenic and disease-linked Tau and GluN2B variants impair homeostatic synaptic scaling of NMDA receptors. (A) Tau KO hippocampal neurons expressing GFP-tagged GluN2B-wild-type with either hTau-Flag wild-type, AT8E, MTBRE or P301L mutants were stimulated with TTX for 24 h. Live neurons were incubated with a rabbit anti-GFP antibody at room temperature to label surface GFP-tagged GluN2B, followed by fixation, permeabilisation, and staining with a mouse anti-GFP antibody to detect total GFP expression. Representative images of the surface and total GFP staining in neurons from each group are shown (top panels). Scale bar, 20 μm. The bottom panels show enlarged images of dendritic segments within the boxed regions. Scale bar, 5 μm. (B) Quantification of the surface/total GFP ratio normalised to that of non-stimulated neurons (n = 18– 36 neurons per group from 3-5 independent experiments). * P < 0.05, ** P < 0.01, **** P < 0.0001 using two-way ANOVA with a Tukey’s multiple comparisons test. All data represent mean ± SEM. (C) Primary hippocampal neurons expressing GFP-tagged GluN2B (wild-type, S1452F, P1439A, L1424F, M1339V, M1331I, K1293R, K1292R, T1272K, A1267S, R1241Q, T1228M). Live neurons were incubated with a rabbit anti-GFP antibody at room temperature to label surface GFP-tagged GluN2B, followed by fixation, permeabilisation, and staining with a mouse anti-GFP antibody to detect total GFP expression. Representative enlarged images of dendritic segments showing the surface and total GFP staining for each group. Scale bar, 5 μm. (D) Quantification of the surface/total GFP ratio in non-stimulated neurons normalised to neurons expressing GFP-tagged GluN2B wild-type (n = 16–39 neurons per group from 2-6 independent experiments). * P < 0.05, ** P < 0.01, **** P < 0.0001 using two-way ANOVA with a Dunnett’s multiple comparisons test. All data represent mean ± SEM. (E) Quantification of the surface/total GFP ratio normalised to the unstimulated group (n = 9–36 neurons per group from 2-6 independent experiments). * P < 0.05, ** P < 0.01, *** P < 0.001, **** P < 0.0001 using an unpaired Mann-Whitney t test. All data represent mean ± SEM.

Disease-associated *GRIN2B* variants are commonly associated with autism spectrum disorders, schizophrenia, epilepsy and intellectual disability (XiangWei et al., 2018). To determine the effects of *GRIN2B* variants on the homeostasis of NMDA receptors in neurons, we examined eleven disease-associated missense variants within the carboxyl-terminal tail of the GluN2B subunit. Under basal conditions, only 3 out of 11 variants (P1439A, K1293R, and R1241Q) exhibited significantly higher surface expression of GFP-GluN2B compared to GFP-GluN2B WT (Figure 3C and 3D). In contrast, all *GRIN2B* variants, except for L1424F, failed to undergo a compensatory increase in surface NMDA receptor expression upon chronic neuronal silencing (Figure 3C and 3E). Our findings suggest that the impairment of homeostatic synaptic scaling of NMDA receptors is a common compensatory mechanism that is dysregulated in many neurodevelopmental disorders and neuropsychiatric diseases.

## DISCUSSION

Here, we report a physiological (non-pathological) role of Tau in mediating the homeostasis of NMDA receptor surface expression. During prolonged neuronal silencing, the phosphorylation of GluN2B on Tyr-1472 is augmented, which prevents its association with AP-2, thereby inhibiting the internalisation of NMDA receptors and enhancing their expression on the neuronal plasma membrane (Lavezzari et al., 2003; Prybylowski et al., 2005; Jang et al., 2015). We show that TTX-induced increase in GluN2B Tyr-1472 phosphorylation and the surface expression of GluN2B-NMDA receptors are prevented in Tau knockout neurons. The phosphorylation of Tyr-1472 is necessary and sufficient to enhance the number of surface GluN2B-NMDA receptors in Tau knockout neurons. Although Tau protein becomes enriched in the postsynaptic compartment to retain active Fyn following TTX treatment, its overall protein level is downregulated. Our data suggest a mechanism that involves a redistribution of Tau molecules into the postsynaptic site and, at the same time, prevents the excessive levels of otherwise normal Tau from neuronal hyperexcitability (Chang et al., 2021b; Chang et al., 2021a; Shao et al., 2022). We propose that the precise control of Tau expression and subcellular localisation is critical for neuronal homeostasis.

The pathological aspect of Tau phosphorylation has been a predominant focus, with a lesser emphasis on its physiological functions at the postsynapse. The site-specific phosphorylation of Tau is tightly regulated to discriminate between pathological and non-pathological conditions. The phosphorylation sites of Tau contributing to disease states have been mapped out and characterised (Chen et al., 2004; Neddens et al., 2018), however, under a non-pathological state, a different set of phosphorylation sites could be necessary for the proper functioning of the Tau protein. This has yet to be characterised thoroughly. Our study reveals two site-specific increases in the phosphorylation of Tau on Ser-235 and Ser-396 upon prolonged neuronal inactivity. Furthermore, we show that the phosphorylation of Ser-235 enhances its interaction with Fyn, due to its proximity to the Fyn binding site (Lee et al., 1998; Lau et al., 2016), and is required for TTX-induced synaptic upscaling of NMDA receptors in neurons. While the Ser-235 phosphorylation site is part of the AT180 Tau epitope, no increase in Thr-231 phosphorylation was observed following TTX treatment. Similarly, the phosphorylation of Ser-396 but not Ser-404 increased during TTX-induced synaptic homeostasis. Both Ser-396 and Ser-404 hyperphosphorylation sites encompass Tau’s paired helical filaments (PHF)-1 epitope. The physiological phosphorylation of Ser-396 but not Ser-404 of Tau have also been reported previously to be required for PICK1-mediated AMPA receptor internalisation during LTD (Regan et al., 2015). These findings suggest that the phosphorylation landscape of Tau is highly site-specific and tightly regulated, enabling the protein to distinguish between physiological and pathological conditions and to facilitate synaptic plasticity.

Finally, we demonstrate that both disease-associated Tau mutations and human GluN2B variants lead to aberrant surface expression of GluN2B-NMDA receptors and impair the neuron’s ability to undergo further compensatory upscaling in response to chronic neuronal inactivity. The heightened expression of GluN2B on the surface correlates with the overactivation of GluN2B, resulting in the excitotoxicity phenotype associated with neurological disease. It is tempting to speculate that the elevated levels of GluN2B-NMDARs on the neuronal surface may not be inherently pathological; it could instead reflect a compensatory response to synaptic weakening, loss or dysfunction, all of which are hallmarks of early neurodegenerative diseases (Styr and Slutsky, 2018).

## MATERIALS AND METHODS

### Animals

Adult female C57BL/6 WT and Tau knockout mice (Tucker et al., 2001) and their embryos (males and females, embryonic day 17) were used to prepare primary hippocampal neurons. All research procedures involving animal use were conducted in accordance with the Australian Code of Practice for the Care and Use of Animals for Scientific Purposes and were approved by the University of Queensland Animal Ethics Committee (2021/AE517).

### HEK293T cells

HEK293T cells were obtained from ATCC (CRL-3216). Cells were grown in Dulbecco’s modified Eagle medium (DMEM) with 4.5g/L glucose (GIBCO) supplemented with 10% fetal bovine serum (FBS, Invitrogen) and 50 U/ml penicillin, 50 μg/ml streptomycin (GIBCO) in a humidified 5% CO_2_ incubator at 37°C.

### DNA constructs

The plasmid encoding GFP-GluN2B was created by subcloning rat GluN2B cDNA into the pRK5 vector and subsequently inserting GFP cDNA after the signal peptide (residues 1-90). GluN2B Y1472E, Y1472F, rare human variants and Tau point mutations were generated with an overlapping PCR protocol. Tau mutant constructs were generated using a FLAG-tagged vector containing human 0N4R TAU (hTAU24, 383aa) as a template. The following expression vectors were generated: R5H, P301L, S235A, S235E; mutant of the AT8 epitope (S202E/T205E/S208E); mutant of the 12E8 epitope (S262E/S356E) and mutants of the AT180 epitope (T231A/S235A and T231E/S235E). Full-length human Fyn and constitutively active Y531F were subcloned from the pcDNA3.1 vector to an FUW-myc lentiviral vector using the unique EcoR*I* and BamH*I* sites. The sgRNA targeting the mouse Tau (5’-GACACAATGGAAGACCATGC-3’) and Fyn (5’-ACTGGGACCCTACGCACGAG-3’) sequence was cloned into the pLentiCRISPR-GFP vector.

### Antibodies

The following antibodies were purchased from commercial sources: mouse anti-β-actin (Cat# sc-47778, Santa Cruz Biotechnology), chicken anti-GFP (Cat# GFP-1020, Aves Labs), mouse anti-GFP (Cat# ab1218, Abcam), mouse anti-myc (Cat# MCA2200, Bio-Rad), rabbit anti-GluN2A (Cat# 04-901, Millipore), rabbit anti-GluN2B (Cat# 06-600, Millipore), rabbit anti-GluN2B phospho-Tyr1472 (Cat# p1516-1472, PhosphoSolution), rabbit anti-GluN2B phospho-S1480 (Cat# PA1-4733, Thermo Fisher Scientific), rabbit anti-Fyn (Cat# 4023, Cell Signaling Technology), mouse anti-Tau (Cat# MAB361, Millipore), mouse anti-Flag (M2, Cat# 1804, Sigma) and rabbit anti-Flag (Cat# 14793, Cell Signaling Technology). Alexa-conjugated and HRP-conjugated secondary antibodies were purchased from Thermo Scientific and Sigma-Aldrich, respectively.

### Rat primary neuronal culture and transient transfection

Hippocampi derived from embryonic day 17 mouse pups were used to prepare primary hippocampal neurons (Yong et al., 2021). Hippocampi were isolated and dissociated with 30 U of papain suspension (Worthington, Lakewood, NJ) for 20 min in a 37°C water bath. A single-cell suspension was obtained by triturating tissues with a P1000 pipette tip and then plated at a density of 12 × 104 cells (per well or 35mm dishes) or 1.2 × 10^6^ cells (per dish) on poly-L-lysine-coated 12-well plates, 6-cm dishes or 35mm MatTek glass bottom dishes in Neurobasal growth medium supplemented with 2 mM Glutamax, 1% penicillin/streptomycin, and 2% B27. Hippocampal neurons were maintained in a Neurobasal medium and fed twice a week. Neurons were kept in a humidified 5% CO_2_ tissue culture incubator at 37°C. Transfection was carried out at DIV 13 with Lipofectamine 2000 (Invitrogen) according to the manufacturer’s instructions. To induce homeostatic plasticity, neurons were treated with TTX (Abcam, 2 μM) at DIV 15 for 24 h to induce synaptic scaling. Cells were processed or imaged at DIV 16.

### Lentivirus packaging and transduction

Lentiviral particles were generated in HEK293T cells that had been transfected with 7 μg of the plasmid of interest, 3 μg each of pMD2.G envelope plasmid, pRSV-Rev encoding plasmid and pMDLg/pRRE packaging constructs via the calcium-phosphate precipitation method. Lentivirus-containing supernatant was collected 48 h after transfection and filtered through a 0.45 μm low protein-binding cellulose acetate membrane. Lentiviral particles were pelleted either by ultracentrifugation at 106,559 *g* on a Beckman SW 32 Ti rotor for 2 h at 4°C or using the PEG-it Virus Precipitation Solution (System Biosciences) following the manufacturer’s protocol. The viral concentrate was resuspended in Neurobasal medium, snap-frozen in liquid nitrogen and stored at −80°C. Hippocampal neurons were transduced with lentiviral particles at DIV 8-9 (overnight) and DIV 12 for sgRNA-knockdown and overexpression, respectively. Transduced neuronal cultures were cultured for a further 3-6 days prior to processing.

### Surface biotinylation assay

The surface biotinylation assay was carried out at DIV 16 to measure the amount of endogenous GluN2A, GluN2B, GluN2B phospho-Y1472 and phospho-S1480 protein on the plasma membrane (Yong et al., 2020; Yong et al., 2021). Live neurons were washed once in ice-cold ACSF (120 mM NaCl, 2 mM CaCl2, 5 mM KCl, 1 mM MgCl2, 30 mM glucose, 25 mM HEPES, pH 8.2), followed by a 30 min incubation with 0.5 mg/ml of sulfo-NHS-SS-biotin (CovaChem) at 4°C. Free biotin was quenched by washing the cells three times with ice-cold TBS (50 mM Tris-HCl, 150 mM NaCl, pH 7.4). Neurons were lysed in RIPA buffer (1% Triton X-100, 0.5% Na-deoxycholate, 0.1% SDS, 2 mM EDTA, 2 mM EGTA, 50 mM NaF, 10 mM Na-pyrophosphate in TBS, pH 7.4) supplemented with EDTA-free protease inhibitor cocktail (Sigma), phosphatase inhibitor cocktail (AG Scientific) and sodium orthovanadate (New England Biology) at 4°C. Lysates were centrifuged at 20,627*g* for 20 min at 4°C and incubated with Neutravidin agarose beads (Thermo Scientific) overnight at 4°C. Beads were washed three times with ice-cold RIPA buffer, and bound proteins were eluted with 2X SDS sample buffer at 50°C for 30 min and analysed by western blotting. The absence of β-actin on the surface fraction confirmed the assay’s specificity.

### Immunolabeling of surface GFP-GluN2B

To determine the levels of GluN2B on the plasma membrane, primary hippocampal neurons transfected with various GFP-GluN2B constructs were incubated with mouse anti-GFP antibody (Abcam, 1:250) for 30 min at 4°C before 10 min fixation in ice-cold parafix solution (4% paraformaldehyde, 4% sucrose in PBS). Neurons were permeabilised (0.25% Triton X-100 in PBS) for 10 min and blocked (10% normal goat serum) for 1 h at room temperature. Total GFP-GluN2B was labelled with chicken anti-GFP antibody (Abcam, 1:5,000) at 4°C overnight. The surface and total GFP-GluN2B were visualised by staining with Alexa-568-conjugated anti-mouse and Alexa-488-conjugated anti-chicken secondary antibodies, respectively. Images were collected with a 63X oil-immersion objective on a Zeiss LSM710 confocal microscope. Fluorescence intensities were quantified using Image J software (National Institutes of Health) for surface and total GluN2B. Data were expressed as the surface/total GluN2B ratio.

### Immunoprecipitation assay

HEK293T cells were co-transfected with either human WT-Fyn-myc or Y531F-Fyn-myc in combination with other human Tau mutant constructs by the calcium phosphate precipitation method. Cells were lysed 24 h later with ice-cold cell lysis buffer (1% Triton X-100, 1mM EDTA, 1 mM EGTA, 50 mM NaF, 5 mM Na-pyrophosphate in PBS) supplemented with EDTA-free protease inhibitor cocktail. Lysates were centrifuged at 20,627*g* for 20 min at 4°C and cleared with protein G-Sepharose beads (Thermo Scientific) for 1 h. Precleared lysates were incubated with protein G-Sepharose beads coupled with antibodies overnight at 4°C. Beads were washed three times with ice-cold cell lysis buffer, and bound proteins were eluted with 2X SDS sample buffer at 100°C for 10 min. Eluted proteins were resolved by SDS-PAGE and analysed by western blotting. Blots were probed using the enhanced chemiluminescence method. Images were acquired on the Odyssey Fc imaging system (LI-COR), and band intensities were quantified using Image Studio Lite software (LI-COR).

### Proximity ligation assay (PLA)

Following TTX-induced homeostatic scaling, the interaction between TAU-Flag (wildtype, S235A phospho-dead and S235E phospho-mimetic) and endogenous Fyn was assessed using the Duolink® In Situ PLA kit (Sigma), according to the manufacturer’s protocol and as previously described (Tan et al., 2023). Neurons were fixed, permeabilised, and blocked before being incubated overnight at room temperature with rabbit anti-Fyn (1:500) and mouse anti-Flag (1:500) primary antibodies and washed with buffer A. PLA probes (mouse PLUS and rabbit MINUS) were applied and incubated at 37 °C for 1 h. The ligation step was carried out for 30 min at 37 °C, followed by two washes. Rolling circle amplification with fluorophore-labelled probe binding was conducted for 100 min at 37 °C. Samples were washed in buffer B and subsequently immunostained with Flag, followed by Alexa Fluor 488-conjugated anti-mouse secondary antibody to visualise neurons expressing TAU-Flag. Neurons were washed and mounted. PLA signals were quantified as the number of puncta across all dendrites, normalised to the region of interest (ROI) area and expressed relative to untreated conditions.

### Immunocytochemistry

Primary hippocampal neurons were washed and fixed with Parafix solution, permeabilized and blocked (10% normal goat serum in PBS) for 1 h. After washing, neurons were incubated with primary antibodies diluted in Can Get Signal® Immunoreaction Enhancer Solution (Cat# NKB-601, Toyobo Life Science) overnight at room temperature. Protein localization was visualized by staining with Alexa-conjugated secondary antibodies for 1 h. Images were collected with a 63X oil-immersion objective on a Zeiss LSM510 confocal microscope. To quantify the colocalization of Copine-6 with intracellular endosomal markers, a region of interest was drawn on the soma to obtain the Mander’s coefficient and Pearson’s correlation coefficient using Just Another Colocalization Plugin (JACoP) in Image J software.

### Subcellular fractionation

Neurons were lysed with ice-cold isotonic buffer (0.32 M sucrose, 4 mM HEPES, pH 7.4) supplemented with an EDTA-free protease inhibitor cocktail, phosphatase inhibitor cocktail, and sodium orthovanadate. Lysates were passed through a 25-gauge needle, and an aliquot was taken for further analysis as the total protein fraction. The remainder of the homogenate was centrifuged at 800*g* for 10 min at 4°C. The supernatant (S1 fraction) was centrifuged for a further 20 min, 10,000*g* at 4°C, and an aliquot of the resulting supernatant was taken for further analysis as the cytosolic protein fraction. The remaining pellet was resuspended in 1 ml of ice-cold isotonic buffer and centrifuged at 10,000*g* for 15 min at 4°C to obtain a crude synaptosomal (P2) fraction. All fractions were denatured and analysed by western blotting.

### Western blotting

Samples were loaded in 7.5% or 12% SDS-PAGE gels and separated at 110 V for 1-2 h. Proteins were then transferred to a PVDF membrane at 100 V for 2 h. Membranes were blocked in 5% skim milk (in TBS containing 0.1% Tween-20, TBS-T) for 1 h and washed in TBS-T three times at 5 min intervals prior to overnight incubation with primary antibodies at 4°C. Membranes were washed in 1% milk/TBS-T five times and incubated with HRP-conjugated secondary antibodies (GE Healthcare, 1:10,000) for 1 h at room temperature. They were washed extensively and developed using the enhanced chemiluminescent method (PerkinElmer). Images were acquired with a LiCOR imaging system and quantified with ImageStudio software.

### Quantification and Statistical Analysis

The sample size (*n*) reported in figure legends represents individual neurons or wells generated from at least three independent experiments, unless otherwise stated. Statistical analysis was performed in Graph Pad Prism 9.0 using one-way analysis of variance (ANOVA) with Tukey’s or Sidak’s post-hoc multiple comparisons tests. For comparison between two groups, a two-tailed unpaired t-test was employed. All data are reported as mean ± standard error of the mean (SEM).

## ACKNOWLEDGEMENTS

This work was supported by Australian National Health and Medical Research Council (NHMRC) Ideas Grant GNT2029692 (to V.A. and X.L.H.Y.) and a generous donation from Kay Bryan OAM. V.A. holds an ARC Future Fellowship (FT220100485). X.L.H.Y. was supported by a Research Training Program Scholarship from the Australian Government, as well as the Ian Lindenmayer PhD Top-up Scholarship. Imaging was performed at the Queensland Brain Institute’s Advanced Microscopy Facility, supported by the Australian Government through ARC LIEF grant LE130100078.

## AUTHOR CONTRIBUTIONS

Conceptualisation: X.L.H.Y and V.A. Methodology: X.L.H.Y., K.S., W.C.K., N.D., A.I., and V.A. Investigation: X.L.H.Y., K.S., W.C.K., I.I., P.P., E.S., A.K., N.D., A.I., J.G., and V.A. Visualisation: X.L.H.Y. and V.A. Funding acquisition: X.L.H.Y. and V.A. Project administration: V.A. Supervision: N.D., A.I., J.G. and V.A. Writing – original draft: X.L.H.Y. and V.A.

---

**Supplementary Figure 1.**
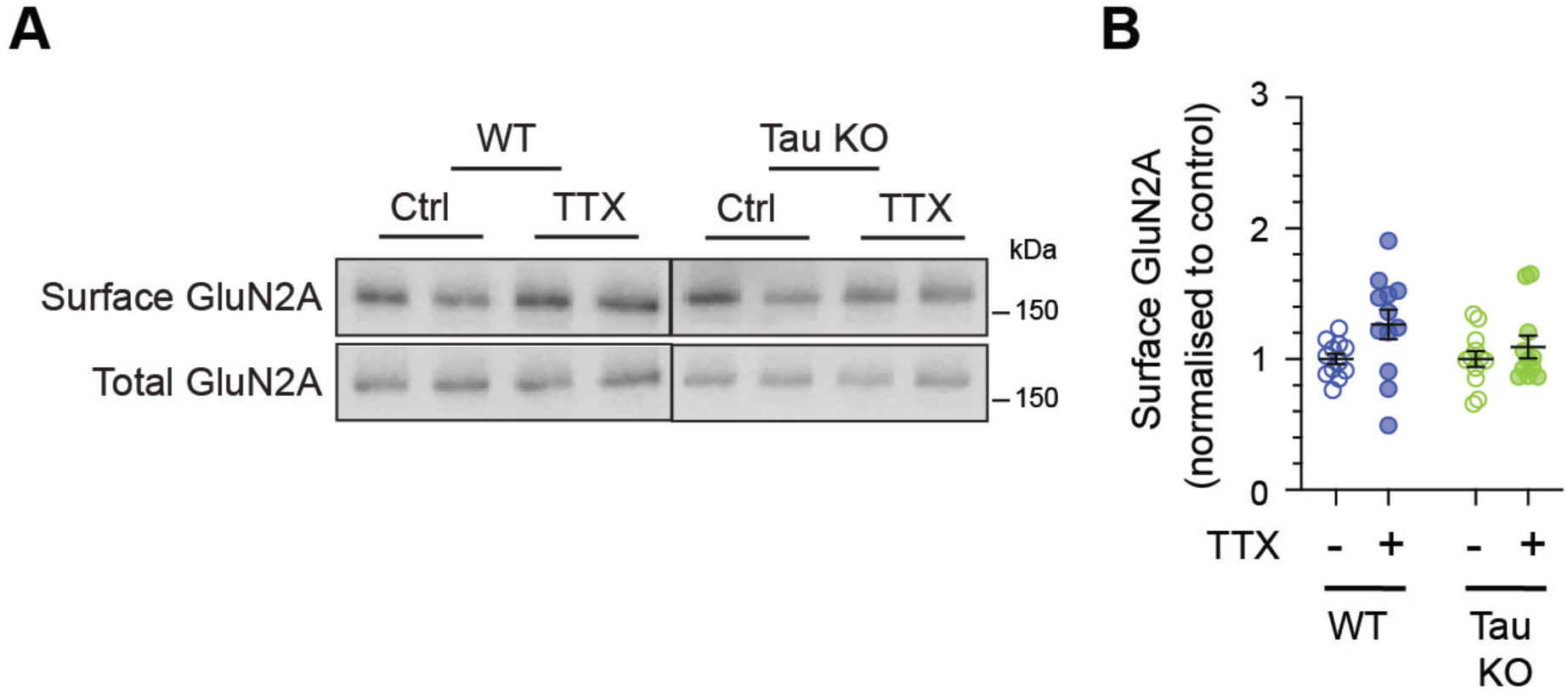
Chronic neuronal inactivity does not upregulate the surface expression of GluN2A-containing NMDA receptors. (A) Primary hippocampal neurons derived from wild-type and Tau knockout mice were treated with water (control) or 2 μM tetrodotoxin (TTX) for 24 h. The relative amounts of surface and total proteins were assessed by western blotting using specific antibodies against GluN2A. This panel was generated using the same samples derived from Figure 1A. (B) Quantification of the surface GluN2B, normalised to the control group (n = 12 cultures per group from 3 independent experiments, using the same samples derived from Figure 1A-C).

**Supplementary Figure 2.**
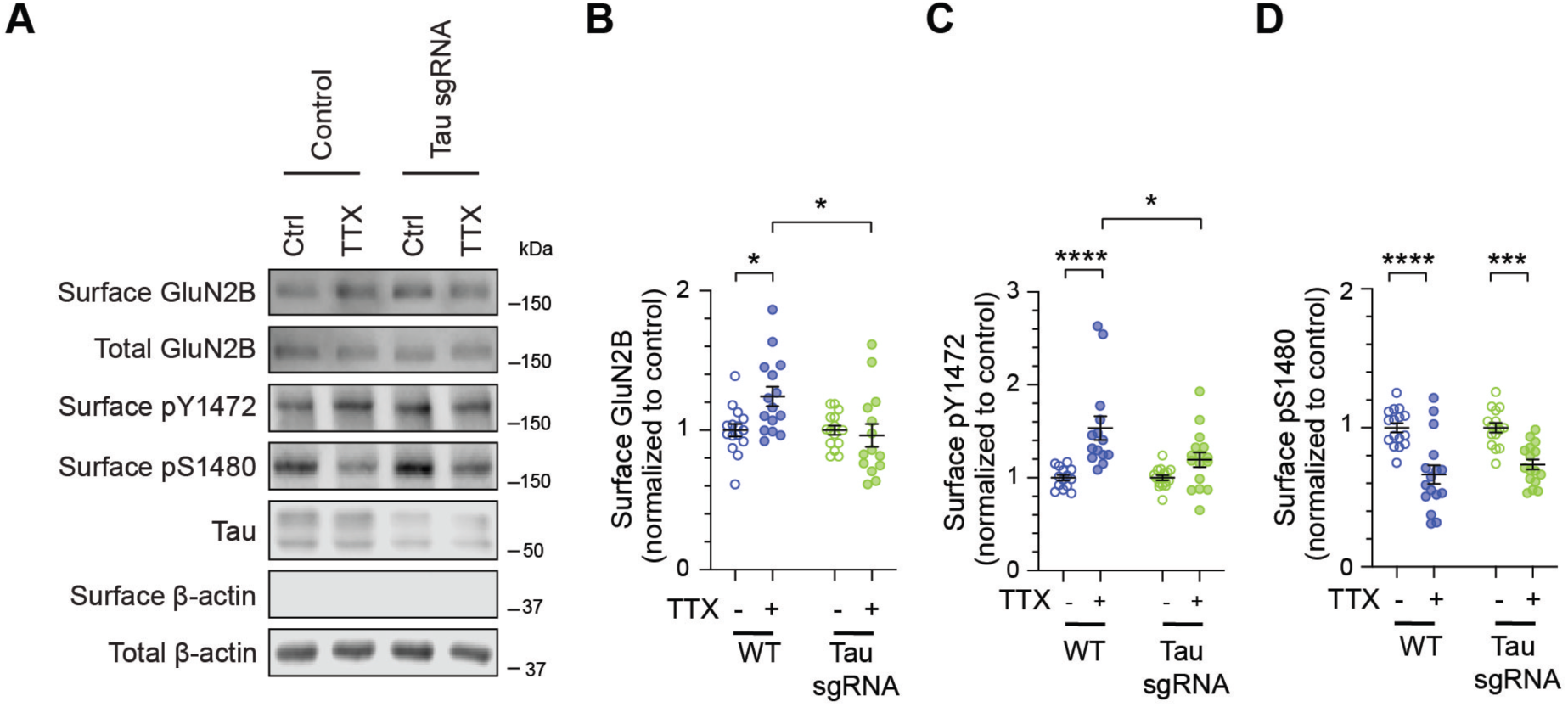
sgRNA-mediated Tau knockdown impairs TTX-induced synaptic upscaling of NMDA receptors. (A) Primary hippocampal neurons were transduced with lentiviral particles expressing Cas9-GFP and Tau sgRNA, at DIV9. Water control or TTX-treated neurons were subjected to a surface biotinylation assay after 24 h treatment. The relative amounts of surface and total proteins were assessed by immunoblotting with specific antibodies against phosphorylated Tyr-1472, Ser-1480, GluN2B, Tau, and β-actin. (B-D) Quantification of the surface GluN2B levels (B), Tyr-1472 phosphorylation (C) and Ser-1480 phosphorylation (D) levels normalised to unstimulated groups (n = 14–15 cultures per group from 4 independent experiments). * P < 0.05, *** P < 0.001, **** P < 0.0001 using two-way ANOVA with a Tukey’s multiple comparison test. All data represent mean ± SEM.

**Supplementary Figure 3.**
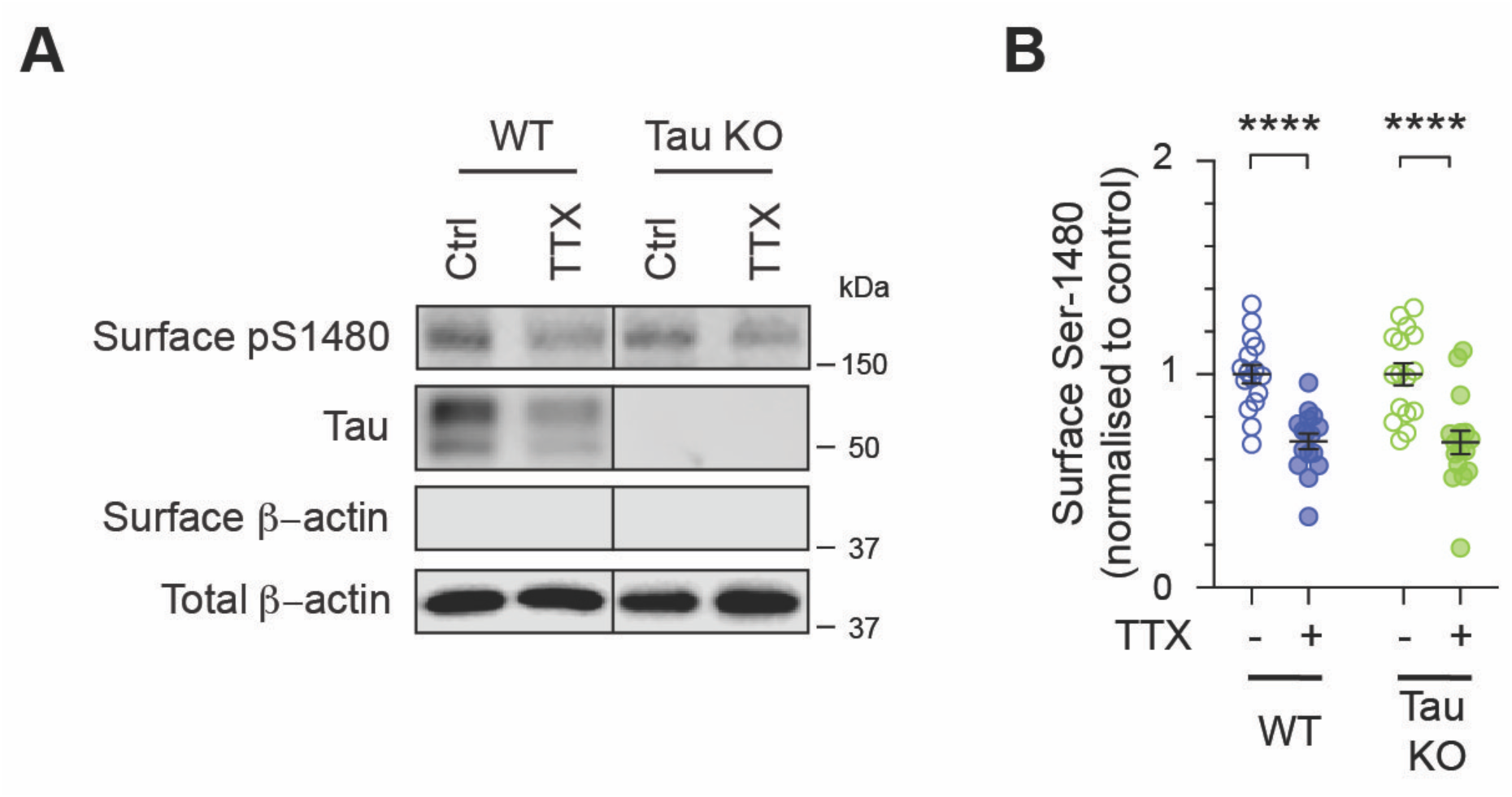
TTX-induced dephosphorylation of GluN2B on Ser-1480 is independent of Tau. (A) Primary hippocampal neurons derived from wild-type and Tau knockout mice were treated with water (control) or 2 μM tetrodotoxin (TTX) for 24 h. The relative amounts of surface and total proteins were assessed by western blotting using specific antibodies against phosphorylated Ser-1480, Tau and β-actin. Blots for β-actin in the Tau KO group were previously shown in Figure 1D, as the same lysates were used. (B) Quantification of the surface GluN2B, normalised to the control group (n = 12 cultures per group from 3 independent experiments, using the same samples derived from Figure 1A-C). **** P < 0.0001 using two-way ANOVA with a Tukey’s multiple comparison test. All data represent mean ± SEM.

**Supplementary Figure 4.**
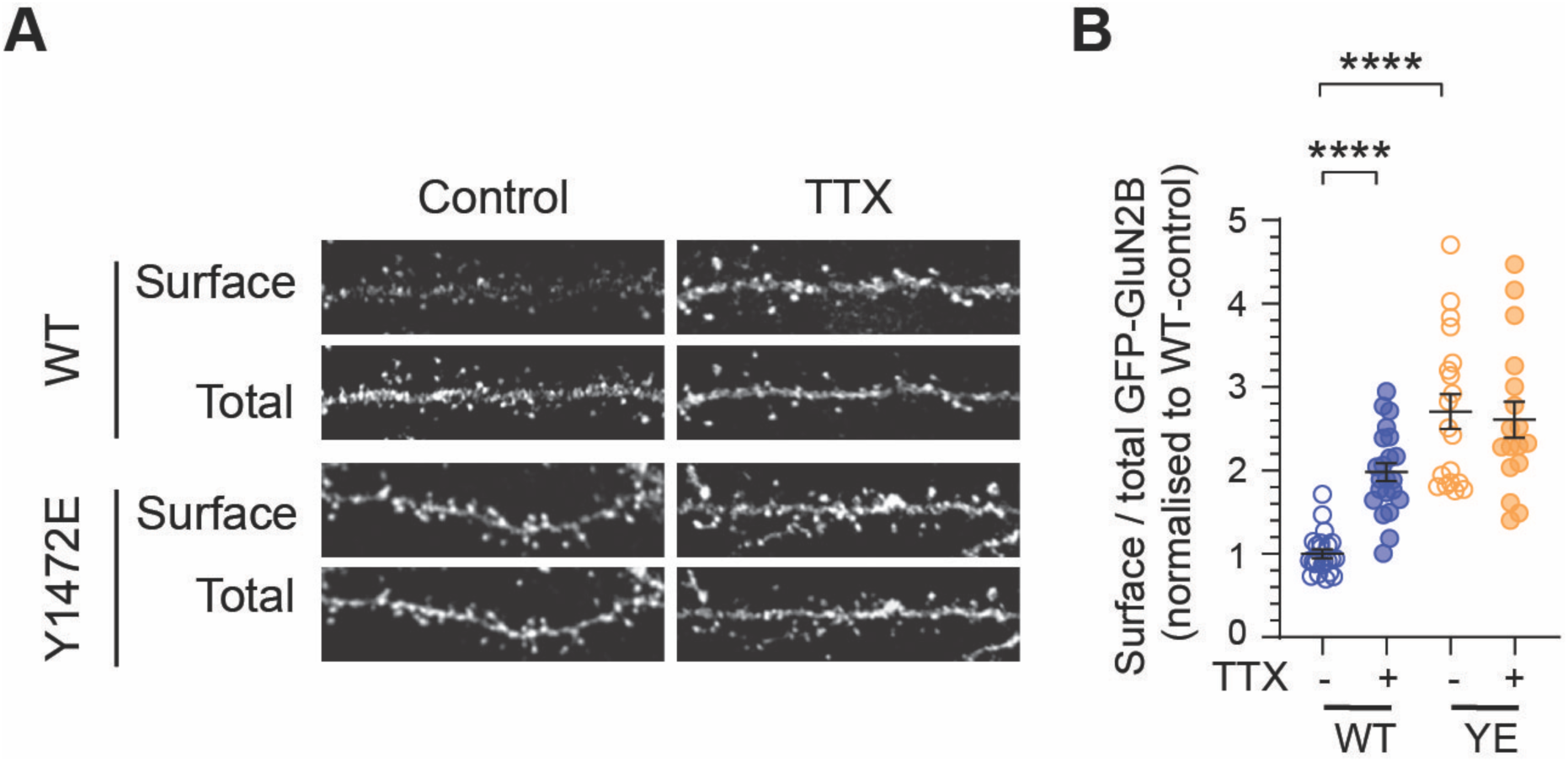
GluN2B pseudo-phosphorylation on Tyr-1472 is sufficient to increase the surface expression of NMDA receptors in Tau knockout neurons. (A) Wild-type hippocampal neurons expressing either GFP-tagged GluN2B-wild-type or Tyr-1472E phospho-mimetic mutant were stimulated with TTX for 24 h. Live neurons were incubated with a rabbit anti-GFP antibody at room temperature to label surface GFP-tagged GluN2B, followed by fixation, permeabilisation, and staining with a mouse anti-GFP antibody to detect total GFP expression. Representative enlarged images of dendritic segments showing the surface and total GFP staining for each group. Scale bar, 5 μm. (B) Quantification of the surface/total GFP ratio normalised to that of non-stimulated neurons (n = 17– 23 neurons per group from 2 independent experiments). **** P < 0.0001 using two-way ANOVA with a Tukey’s multiple comparison test. All data represent mean ± SEM.

**Supplementary Figure 5.**
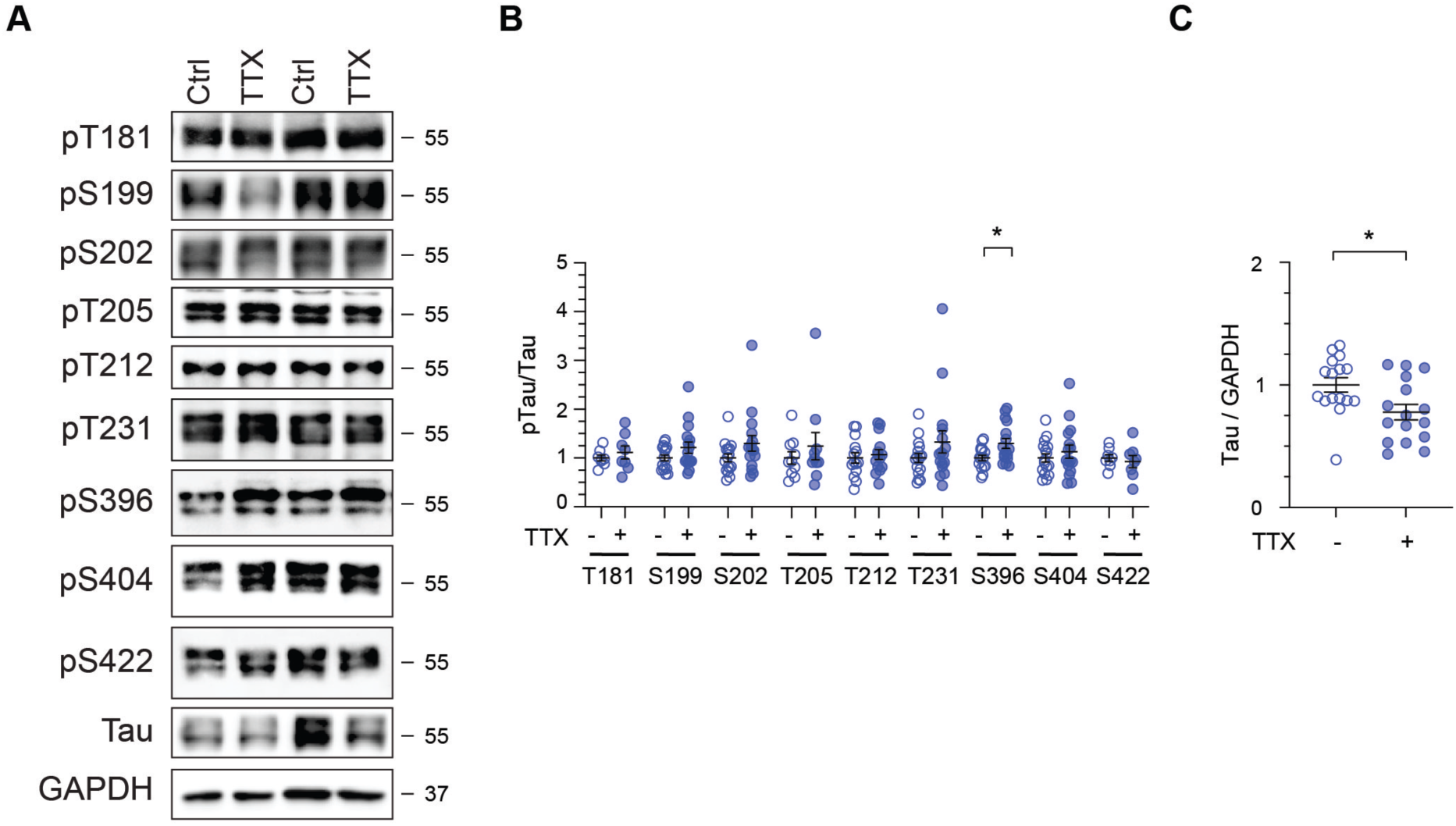
Site-specific phosphorylation of Tau in response to prolonged neuronal silencing. (A) Total sample lysates were prepared from wild-type hippocampal neurons following 24 h water control or TTX treatment. The relative amounts of proteins were assessed by western blotting using antibodies specific for phosphorylated Tau, total Tau, and GAPDH. Blots for Tau and GAPDH were previously shown in Figure 2A, as the same lysates were used. (B) Quantification of the ratios of phosphorylated Tau/total Tau. * P < 0.05 using an unpaired Mann-Whitney t test. All data are represented as mean ± SEM. (C) Quantification of the ratios of phosphorylated Tau/GAPDH (n = 16 cultures per group from 4 independent experiments). * P < 0.05 using an unpaired Mann-Whitney t test. All data are represented as mean ± SEM.

## REFERENCES

Anggono V, Clem RL, Huganir RL (2011) PICK1 loss of function occludes homeostatic synaptic scaling. J Neurosci 31:2188–2196.

Bateup HS, Denefrio CL, Johnson CA, Saulnier JL, Sabatini BL (2013) Temporal dynamics of a homeostatic pathway controlling neural network activity. Front Mol Neurosci 6:28.

Bi M, Gladbach A, van Eersel J, Ittner A, Przybyla M, van Hummel A, Chua SW, van der Hoven J, Lee WS, Muller J, Parmar J, Jonquieres GV, Stefen H, Guccione E, Fath T, Housley GD, Klugmann M, Ke YD, Ittner LM (2017) Tau exacerbates excitotoxic brain damage in an animal model of stroke. Nat Commun 8:473.

Brigman JL, Wright T, Talani G, Prasad-Mulcare S, Jinde S, Seabold GK, Mathur P, Davis MI, Bock R, Gustin RM, Colbran RJ, Alvarez VA, Nakazawa K, Delpire E, Lovinger DM, Holmes A (2010) Loss of GluN2B-containing NMDA receptors in CA1 hippocampus and cortex impairs long-term depression, reduces dendritic spine density, and disrupts learning. J Neurosci 30:4590–4600.

Caya-Bissonnette L, Béïque JC (2024) Half a century legacy of long-term potentiation. Curr Biol 34:R640–R662.

Chang CW, Shao E, Mucke L (2021a) Tau: Enabler of diverse brain disorders and target of rapidly evolving therapeutic strategies. Science 371.

Chang CW, Evans MD, Yu X, Yu GQ, Mucke L (2021b) Tau reduction affects excitatory and inhibitory neurons differently, reduces excitation/inhibition ratios, and counteracts network hypersynchrony. Cell Rep 37:109855.

Chen F, David D, Ferrari A, Gotz J (2004) Posttranslational modifications of tau--role in human tauopathies and modeling in transgenic animals. Curr Drug Targets 5:503–515.

Chung HJ, Huang YH, Lau LF, Huganir RL (2004) Regulation of the NMDA receptor complex and trafficking by activity-dependent phosphorylation of the NR2B subunit PDZ ligand. J Neurosci 24:10248–10259.

Davis GW, Bezprozvanny I (2001) Maintaining the stability of neural function: a homeostatic hypothesis. Annu Rev Physiol 63:847–869.

Ehlers MD (2003) Activity level controls postsynaptic composition and signaling via the ubiquitin-proteasome system. Nat Neurosci 6:231–242.

Fernandes D, Carvalho AL (2016) Mechanisms of homeostatic plasticity in the excitatory synapse. J Neurochem 139:973–996.

Frere S, Slutsky I (2018) Alzheimer’s Disease: From Firing Instability to Homeostasis Network Collapse. Neuron 97:32–58.

Goel A, Jiang B, Xu LW, Song L, Kirkwood A, Lee HK (2006) Cross-modal regulation of synaptic AMPA receptors in primary sensory cortices by visual experience. Nat Neurosci 9:1001–1003.

Hansen KB, Wollmuth LP, Bowie D, Furukawa H, Menniti FS, Sobolevsky AI, Swanson GT, Swanger SA, Greger IH, Nakagawa T, McBain CJ, Jayaraman V, Low CM, Dell’Acqua ML, Diamond JS, Camp CR, Perszyk RE, Yuan H, Traynelis SF, Barker E (2021) Structure, function, and pharmacology of glutamate receptor ion channels. Pharm Rev 73:298–487.

Hardingham GE, Bading H (2010) Synaptic versus extrasynaptic NMDA receptor signalling: implications for neurodegenerative disorders. Nat Rev Neurosci 11:682–696.

Ittner A, Ittner LM (2018) Dendritic tau in Alzheimer’s disease. Neuron 99:13–27.

Ittner LM, Ke YD, Delerue F, Bi M, Gladbach A, van Eersel J, Wolfing H, Chieng BC, Christie MJ, Napier IA, Eckert A, Staufenbiel M, Hardeman E, Gotz J (2010) Dendritic function of tau mediates amyloid-β toxicity in Alzheimer’s disease mouse models. Cell 142:387–397.

Jang SS, Royston SE, Xu J, Cavaretta JP, Vest MO, Lee KY, Lee S, Jeong HG, Lombroso PJ, Chung HJ (2015) Regulation of STEP61 and tyrosine-phosphorylation of NMDA and AMPA receptors during homeostatic synaptic plasticity. Mol Brain 8:55.

Jumper J et al. (2021) Highly accurate protein structure prediction with AlphaFold. Nature 596:583–589.

Keck T, Keller GB, Jacobsen RI, Eysel UT, Bonhoeffer T, Hubener M (2013) Synaptic scaling and homeostatic plasticity in the mouse visual cortex in vivo. Neuron 80:327–334.

Keck T, Toyoizumi T, Chen L, Doiron B, Feldman DE, Fox K, Gerstner W, Haydon PG, Hubener M, Lee HK, Lisman JE, Rose T, Sengpiel F, Stellwagen D, Stryker MP, Turrigiano GG, van Rossum MC (2017) Integrating Hebbian and homeostatic plasticity: the current state of the field and future research directions. Philos Trans R Soc Lond B Biol Sci 372.

Lau DH, Hogseth M, Phillips EC, O’Neill MJ, Pooler AM, Noble W, Hanger DP (2016) Critical residues involved in tau binding to fyn: implications for tau phosphorylation in Alzheimer’s disease. Acta Neuropathol Commun 4:49.

Lavezzari G, McCallum J, Lee R, Roche KW (2003) Differential binding of the AP-2 adaptor complex and PSD-95 to the C-terminus of the NMDA receptor subunit NR2B regulates surface expression. Neuropharmacology 45:729–737.

Lee G, Newman ST, Gard DL, Band H, Panchamoorthy G (1998) Tau interacts with src-family non-receptor tyrosine kinases. J Cell Sci 111:3167–3177.

Liu F, Iqbal K, Grundke-Iqbal I, Gong CX (2002) Involvement of aberrant glycosylation in phosphorylation of tau by cdk5 and GSK-3beta. FEBS Lett 530:209–214.

Lussier MP, Sanz-Clemente A, Roche KW (2015) Dynamic regulation of N-methyl-d-aspartate (NMDA) and α-amino-3-hydroxy-5-methyl-4-isoxazolepropionic acid (AMPA) receptors by posttranslational modifications. J Biol Chem 290:28596–28603.

Maffei A, Nelson SB, Turrigiano GG (2004) Selective reconfiguration of layer 4 visual cortical circuitry by visual deprivation. Nat Neurosci 7:1353–1359.

Marder E, Prinz AA (2003) Current compensation in neuronal homeostasis. Neuron 37:2–4.

Nakazawa T, Komai S, Tezuka T, Hisatsune C, Umemori H, Semba K, Mishina M, Manabe T, Yamamoto T (2001) Characterization of Fyn-mediated tyrosine phosphorylation sites on GluR epsilon 2 (NR2B) subunit of the N-methyl-D-aspartate receptor. J Biol Chem 276:693–699.

Neddens J, Temmel M, Flunkert S, Kerschbaumer B, Hoeller C, Loeffler T, Niederkofler V, Daum G, Attems J, Hutter-Paier B (2018) Phosphorylation of different tau sites during progression of Alzheimer’s disease. Acta Neuropathol Commun 6:52.

O’Brien RJ, Kamboj S, Ehlers MD, Rosen KR, Fischbach GD, Huganir RL (1998) Activity-dependent modulation of synaptic AMPA receptor accumulation. Neuron 21:1067–1078.

Padmanabhan P, Martinez-Marmol R, Xia D, Gotz J, Meunier FA (2019) Frontotemporal dementia mutant Tau promotes aberrant Fyn nanoclustering in hippocampal dendritic spines. Elife 8:e45040.

Palop JJ, Mucke L (2010) Amyloid-beta-induced neuronal dysfunction in Alzheimer’s disease: from synapses toward neural networks. Nat Neurosci 13:812–818.

Paoletti P, Bellone C, Zhou Q (2013) NMDA receptor subunit diversity: impact on receptor properties, synaptic plasticity and disease. Nat Rev Neurosci 14:383–400.

Parra Bravo C, Naguib SA, Gan L (2024) Cellular and pathological functions of tau. Nat Rev Mol Cell Biol 25:845–864.

Pettersen EF, Goddard TD, Huang CC, Meng EC, Couch GS, Croll TI, Morris JH, Ferrin TE (2021) UCSF ChimeraX: Structure visualization for researchers, educators, and developers. Protein Sci 30:70–82.

Polanco JC, Li C, Bodea LG, Martinez-Marmol R, Meunier FA, Gotz J (2018) Amyloid-β and tau complexity - towards improved biomarkers and targeted therapies. Nat Rev Neurol 14:22–39.

Prybylowski K, Chang K, Sans N, Kan L, Vicini S, Wenthold RJ (2005) The synaptic localization of NR2B-containing NMDA receptors is controlled by interactions with PDZ proteins and AP-2. Neuron 47:845–857.

Regan P, Piers T, Yi JH, Kim DH, Huh S, Park SJ, Ryu JH, Whitcomb DJ, Cho K (2015) Tau phosphorylation at serine 396 residue is required for hippocampal LTD. J Neurosci 35:4804–4812.

Sakaue F, Saito T, Sato Y, Asada A, Ishiguro K, Hasegawa M, Hisanaga S (2005) Phosphorylation of FTDP-17 mutant tau by cyclin-dependent kinase 5 complexed with p35, p25, or p39. J Biol Chem 280:31522–31529.

Sanz-Clemente A, Matta JA, Isaac JT, Roche KW (2010) Casein kinase 2 regulates the NR2 subunit composition of synaptic NMDA receptors. Neuron 67:984–996.

Shao E, Chang CW, Li Z, Yu X, Ho K, Zhang M, Wang X, Simms J, Lo I, Speckart J, Holtzman J, Yu GQ, Roberson ED, Mucke L (2022) TAU ablation in excitatory neurons and postnatal TAU knockdown reduce epilepsy, SUDEP, and autism behaviors in a Dravet syndrome model. Sci Transl Med 14:eabm5527.

Soares C, Lee KF, Nassrallah W, Beique JC (2013) Differential subcellular targeting of glutamate receptor subtypes during homeostatic synaptic plasticity. J Neurosci 33:13547–13559.

Styr B, Slutsky I (2018) Imbalance between firing homeostasis and synaptic plasticity drives early-phase Alzheimer’s disease. Nat Neurosci 21:463–473.

Tan JZA, Jang SE, Batallas-Borja A, Bhembre N, Chandra M, Zhang L, Guo H, Ringuet MT, Widagdo J, Collins BM, Anggono V (2023) Copine-6 is a Ca2+ sensor for activity-induced AMPA receptor exocytosis. Cell Rep 42:113460.

Tucker KL, Meyer M, Barde YA (2001) Neurotrophins are required for nerve growth during development. Nat Neurosci 4:29–37.

Turrigiano GG (1999) Homeostatic plasticity in neuronal networks: the more things change, the more they stay the same. Trends Neurosci 22:221–227.

Turrigiano GG (2008) The self-tuning neuron: synaptic scaling of excitatory synapses. Cell 135:422–435.

Turrigiano GG (2017) The dialectic of Hebb and homeostasis. Philos Trans R Soc Lond B Biol Sci 372. Turrigiano GG, Nelson SB (2004) Homeostatic plasticity in the developing nervous system. Nat Rev Neurosci 5:97–107.

Turrigiano GG, Leslie KR, Desai NS, Rutherford LC, Nelson SB (1998) Activity-dependent scaling of quantal amplitude in neocortical neurons. Nature 391:892–896.

Vieira M, Yong XLH, Roche KW, Anggono V (2020) Regulation of NMDA glutamate receptor functions by the GluN2 subunits. J Neurochem 154:121–143.

von Engelhardt J, Doganci B, Jensen V, Hvalby O, Gongrich C, Taylor A, Barkus C, Sanderson DJ, Rawlins JN, Seeburg PH, Bannerman DM, Monyer H (2008) Contribution of hippocampal and extra-hippocampal NR2B-containing NMDA receptors to performance on spatial learning tasks. Neuron 60:846–860.

Watt AJ, van Rossum MC, MacLeod KM, Nelson SB, Turrigiano GG (2000) Activity coregulates quantal AMPA and NMDA currents at neocortical synapses. Neuron 26:659–670.

Wu CH, Ramos R, Katz DB, Turrigiano GG (2021) Homeostatic synaptic scaling establishes the specificity of an associative memory. Curr Biol 31:2274–2285 e2275.

Wu D, Bacaj T, Morishita W, Goswami D, Arendt KL, Xu W, Chen L, Malenka RC, Sudhof TC (2017) Postsynaptic synaptotagmins mediate AMPA receptor exocytosis during LTP. Nature 544:316–321.

XiangWei W, Jiang Y, Yuan H (2018) De novo mutations and rare variants occurring in NMDA receptors. Curr Opin Physiol 2:27–35.

Yashiro K, Philpot BD (2008) Regulation of NMDA receptor subunit expression and its implications for LTD, LTP, and metaplasticity. Neuropharmacology 55:1081–1094.

Yong XLH, Cousin MA, Anggono V (2020) PICK1 controls activity-dependent synaptic vesicle cargo retrieval. Cell Rep 33:108312.

Yong XLH, Zhang L, Yang L, Chen X, Tan JZA, Yu X, Chandra M, Livingstone E, Widagdo J, Vieira MM, Roche KW, Lynch JW, Keramidas A, Collins BM, Anggono V (2021) Regulation of NMDA receptor trafficking and gating by activity-dependent CaMKIIα phosphorylation of the GluN2A subunit. Cell Rep 36:109338.

